# More partners, more ranges: generalist legumes spread more easily around the globe

**DOI:** 10.1101/406850

**Authors:** Tia L. Harrison, Anna K. Simonsen, John R. Stinchcombe, Megan E. Frederickson

## Abstract

How does mutualism affect range expansion? On one hand, mutualists might thrive in new habitats thanks to the resources, stress tolerance, or defense provided by their partners. On the other, specialized mutualists might fail to find compatible partners beyond their range margins, limiting further spread. A recent global analysis of legume ranges found that non-symbiotic legumes have been successfully introduced to more ranges than legumes that form symbioses with rhizobia, but there is still abundant unexplained variation in introduction success within symbiotic legumes. Here, we test the hypothesis that generalist legumes have spread to more introduced ranges than specialist legumes. We used published data and rhizobial 16S rRNA sequences from GenBank to quantify the number of rhizobia partners that associate with each of 159 legume species, spanning the legume phylogeny and the globe. We found that generalist legumes occur in more introduced ranges than specialist legumes, suggesting that among mutualists, specialization hinders range expansions.

## 1. Introduction

Whether reciprocally beneficial species interactions, or mutualisms, constrain or facilitate range expansions is a fundamental question in evolution and ecology [1]. Mutualism might facilitate range expansion by helping organisms tolerate abiotic stress, overcome resource limitation, or resist enemies in new environments [1-3], but this requires that mutualists find appropriate partners wherever they go. Generalists may be more likely than specialists to acquire mutualist partners in a new range, and thus may spread more easily around the globe. Here, we test the hypothesis that generalist legumes, capable of associating with a greater diversity of bacterial mutualists, have been successfully introduced to more novel ranges than specialist legumes.

Rhizobia fix atmospheric nitrogen for their legume hosts, and consequently the mutualism between legumes and rhizobia can help plants establish in new habitats with nutrient-deficient soils [4]. However, a global analysis of legume introductions [5] found that non-symbiotic legumes have spread to more new ranges than legumes that host rhizobia, suggesting that symbiotic legumes have trouble finding compatible rhizobia in novel environments. Their findings suggest a corollary prediction, which to date remains untested: by the same logic, one would predict that specialist symbiotic legumes would have a harder time finding compatible mutualists, and thus establish in fewer introduced ranges.

Previous work legumes has produced conflicting results on the relationship between rhizobia diversity and introduction success, with some studies finding no consistent difference in rhizobial diversity between invasive and non-invasive legumes [6,7] and others finding that invasive legumes are more likely to be generalists [8,9]. These studies have focused on a single legume clade or performed their experiments or surveys on a regional scale, potentially limiting the generality of their conclusions. We assess the degree of legume specialization on rhizobia across the legume phylogeny to understand how mutualism specificity impacts establishment outside the native range at a global scale. We combine published data on legume-rhizobia associations, rhizobia 16S sequence data from GenBank, and legume range data from Simonsen et al [5] to ask: have generalist legumes been introduced to more new ranges compared to specialist legumes? We find that generalist legume species have indeed established in more non-native ranges than specialist legumes.

## 2. Materials and methods

### Rhizobia richness

We characterized the richness of rhizobial taxa that a legume species associates with using two metrics. First, we summed the number of rhizobia genera associated with 159 legume species in a list compiled by Andrews and Andrews [10]; they assembled a list of rhizobia genera known to associate with a variety of legume genera by searching published literature (Web of Science).

Second, we searched the NCBI GenBank database for 16S rhizobia sequences and clustered the sequences into operational taxonomic units (OTUs). To calculate the number of unique OTUs, we filtered for sequences that had a minimum sequence length of 200 bp, location information (country), and a plant host identified to species. Our analysis incorporated 73 host plant species and we used only the sequences from the native range to calculate OTUs for each host species. We aligned sequences for each plant species using MUSCLE, setting the maxiters parameter to 2, as recommended for large alignments [11]. We clustered aligned 16S sequences in the program Mothur [12] using the furthest neighbouring algorithm and assigned sequences with 97% similarity to a single OTU [12]. We calculated OTUs both with and without sub-sampling (details in Supplementary Materials) for plant species that had a minimum of 10 sequences. The sub-sampled results (data not shown) were qualitatively similar to the results without sub-sampling and so we only report the latter. Although rhizobia 16S sequences on GenBank are likely a non-random sample of rhizobia that associate with legumes, coming primarily from well-studied and often widely distributed legume species, we plotted the number of sequences collected from GenBank per country across the globe (Supplemental Fig. 1) and found fairly good global coverage, with the main exception being an absence of rhizobial sequences from many African countries.

### Comparing introduced and native rhizobia communities

We identified how many rhizobia taxa are shared between the native and introduced range of each legume species by comparing each plant’s rhizobia community in both ranges using unweighted unifrac distance. Therefore, we calculated unifrac scores for all legume species that had at least one introduced range. We coded each legume species as having a native and introduced range that either a) share at least one rhizobia taxon (unifrac score less than 1) or b) share no rhizobia taxa (unifrac score of 1). To calculate unweighted unifrac distance between introduced range and native range rhizobia sequences, we first identified each of the raw 16S sequences used to develop the OTU dataset as belonging to either the plant host’s native or introduced range using the country information for each sequence. We then aligned sequences associated with each plant species using the program MUSCLE [11]. We calculated unweighted unifrac distance between the native and introduced sequences in the program mothur by filtering sequences and building trees using the function clearcut [12].

**Table 1:** Legume introduction success (number of introduced ranges) predicted by rhizobia richness, rhizobia community, plant life history, plant human usage, plant locality, and plant range area.

| Factor | Estimate | Standard error | Df | F | p |
| --- | --- | --- | --- | --- | --- |
| *Genus results, df=149, lambda=0.292* |  |  |  |  |  |
| **Number of genera** | **2.00** | **1.19** | **1** | **19.89** | **<0.0001** |
| Annual | 1.02 | 2.66 | 1 | 1.15 | 0.2853 |
| **Number of human uses** | **4.68** | **0.41** | **1** | **103.99** | **<0.0001** |
| **Latitude of native range** | **-0.10** | **0.05** | **1** | **7.82** | **0.0058** |
| **Area of native range** | **-4.81** | **1.13** | **1** | **18.19** | **<0.0001** |
| *OTU results, df=54, lambda=0.241* |  |  |  |  |  |
| **OTUs (log)** | **12.70** | **5.00** | **1** | **14.76** | **0.0003** |
| **Rhizobia community (shared)** | **7.73** | **3.58** | **1** | **9.60** | **0.0031** |
| Annual | 2.85 | 4.91 | 1 | 2.02 | 0.1610 |
| **Number of human uses** | **2.72** | **0.81** | **1** | **8.03** | **0.0065** |
| Latitude of native range | -0.07 | 0.10 | 1 | 2.14 | 0.1489 |
| **Area of native range** | **-5.04** | **2.07** | **1** | **5.91** | **0.0189** |
| Number of sequences | -0.04 | 0.03 | 1 | 2.26 | 0.1383 |

### Phylogenetic least squares tests

To determine if the number of genera or rhizobia OTUs is associated with legume establishment outside the native range, we fit phylogenetic least squares (PGLS) models with the number of introduced ranges as the response variable and number of partners (genera or OTUs) as the predictor variable. Simonsen et al. [5] calculated the number of ranges from geographic distribution data compiled by the International Legume Database and Information Service (ILDIS) by defining geographic regions with polygons and lumping contiguous polygons for each legume species. Legumes with no introduced ranges were included in the analysis.

We used PGLS models to control for phylogenetic non-independence among the plant species in the dataset using the package caper in R [13] and an angiosperm phylogeny [14] pruned to include only the legume species in our dataset. We log-transformed the number of OTUs because the data were right-skewed. We obtained similar results using the raw number of OTUs (not shown). We included a number of covariates in the analysis including number of human uses, latitude of native range, area of native range, and one plant life history trait (annual or perennial). The area of native range variable was mean-centered and scaled by the standard deviation. These covariates were scraped from the ILDIS and reported by Simonsen et al.[5].

In the OTU PGLS analysis, we included number of sequences as a covariate to account for variation in the number of rhizobia sequences per plant species. Also in the OTU model, we included rhizobia community categories to test whether plants that associate with more phylogenetically related rhizobia communities in their introduced and native range are able to spread to more ranges. We expected that plant species that are able to find similar rhizobia communities outside their native range would be introduced to more ranges. We allowed the parameter λ (Pagel’s λ) to vary and optimize in our model to estimate and account for any phylogenetic signal. We checked model diagnostic plots to ensure that the phylogenetic residuals met the assumptions of the PGLS model. We inspected quantile-quantile plots for linearity, plots of phylogenetic residuals versus fitted values for a random scattering of points, and removed all outliers with phylogenetic residuals greater than ± 3 [13].

## 3. Results and Discussion

### Legumes with more rhizobia partners are introduced to more ranges

We found that legume establishment outside the native range was significantly related to both the number of rhizobia genera and OTUs (Table 1, Figure 1), suggesting that legumes with more rhizobial partners are introduced to more ranges. We observed moderate phylogenetic signal in the introduced range variable in our PGLS model (Genus: λ = 0.292; OTU: λ = 0.339), indicating that introduction success is somewhat phylogenetically conserved. In contrast, Simonsen et al [5] found much weaker phylogenetic signal in their dataset perhaps because they analyzed a much larger dataset.

**Figure 1.**
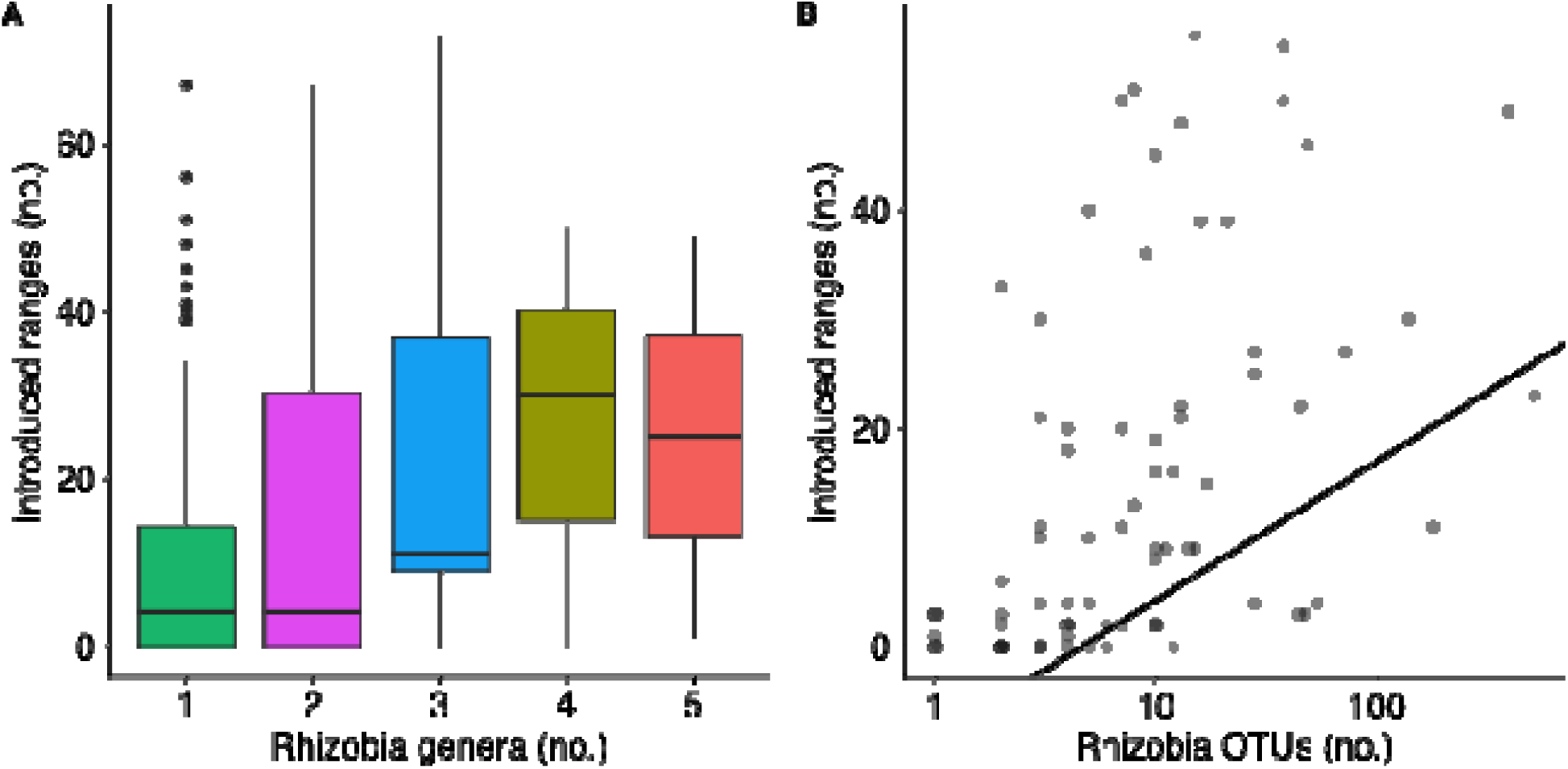
Relationship between introduction success and number of rhizobia A) genera or B) OTUs calculated from native range sequences. Each dot is a legume species. PGLS model estimates are reported in Table 1. Regression line in B) is estimated from the PGLS model.

**Figure 2.**
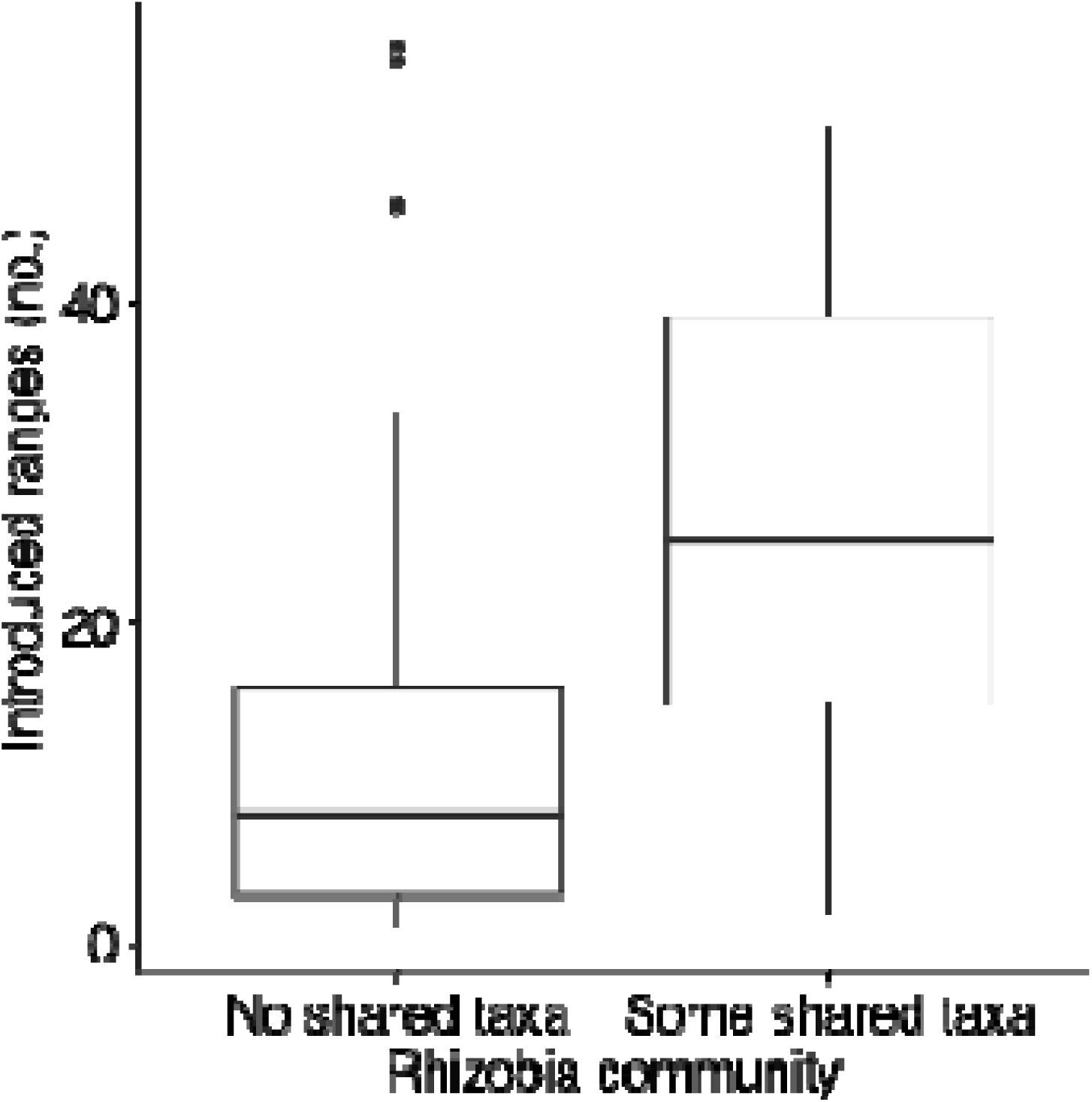
Relationship between introduction success and rhizobia community comparisons between the native and introduced range. PGLS model estimates are reported in Table 1.

Overall, our results suggest that a generalist strategy is important for species across the legume phylogeny in establishing in new ranges around the world. These results support the hypothesis that generalist legumes are able to form mutualisms in many different environments because they can associate with a diversity of rhizobia in their native range (OTU results: Fig. 1B). Specialist legumes are less likely to find a compatible rhizobium partner in novel habitats and thus may fail to establish because they lack mutualists that provide nitrogen. Specialist legumes that have been introduced to a large number of ranges could have been introduced to their new environments together with their compatible rhizobia strain, either by human intervention or by rhizobial contamination of surrounding soil or seeds. We did find that number of human uses was a significant covariate in influencing number of introduced ranges (Table 1) in both our genus level and OTU level results, suggesting that human intervention is a likely explanation for the introduction of specialist legume species.

We also found that rhizobia community composition impacts introduction success in legumes (Table 1: OTU results). In particular, our results show that legumes that share rhizobial taxa in both their native and introduced range have been introduced to more ranges compared to legumes that share no rhizobia taxa (Fig. 3). Our results suggest that legumes that are able to establish in many ranges are able to do so because they find similar and thus compatible rhizobia outside their native range. Legumes species that share no rhizobia taxa between their native and introduced ranges may have spread to fewer new ranges because they cannot form effective mutualisms with the phylogenetically distinct rhizobia taxa in those areas.

### Conclusion

Our study highlights how a generalist strategy can strongly influence the distribution of symbiotic legume species across the globe. Generalist strategies provide many benefits to symbiotic legumes [15] and our study suggests that associating with many rhizobia partners in the native range may increase the probability that a legume will find at least one compatible rhizobia in their introduced range. Diversity in mutualism partners could be an important factor for facilitating range expansions in many other globally widespread mutualisms.

## Data accessibility

A list of legume species and a table of the accession numbers for sequences used in the analysis will be uploaded on Dryad upon acceptance.

## Competing interests

We have no conflict of interest.

## Author contributions

TLH, AKS, JRS, and MEF developed the study and wrote the manuscript. TLH and AKS collected the data. TLH, AKS, JRS, and MEF analyzed and interpreted the data.

### Acknowledgements

We thank Stephen Wright and Corlett Wood for comments on the project. We thank Luke Mahler and Katrina Kaur for assistance with phylogenetic analysis.

## Funding

NSERC Discovery Grant (JRS, MEF)

NSERC Graduate Scholarship (TLS)

